## Supplemental Materials for "Developmental trajectories of EEG aperiodic and periodic components: Implications for understanding thalamocortical development during infancy"

**Supplemental Figure 1 - Electrode layout:** (A) 128-channel Hydrocel Geodesic Sensor Net. (B) 64-channel Geodesic Sensor Net. Pink circles denote 10-20 electrodes, and blue circles denote the additional electrodes included in ICA and MARA steps of pre-processing. (C), (D) Electrodes averaged for frontal (yellow), central (blue), temporal (orange), and posterior (green) regions of interest.

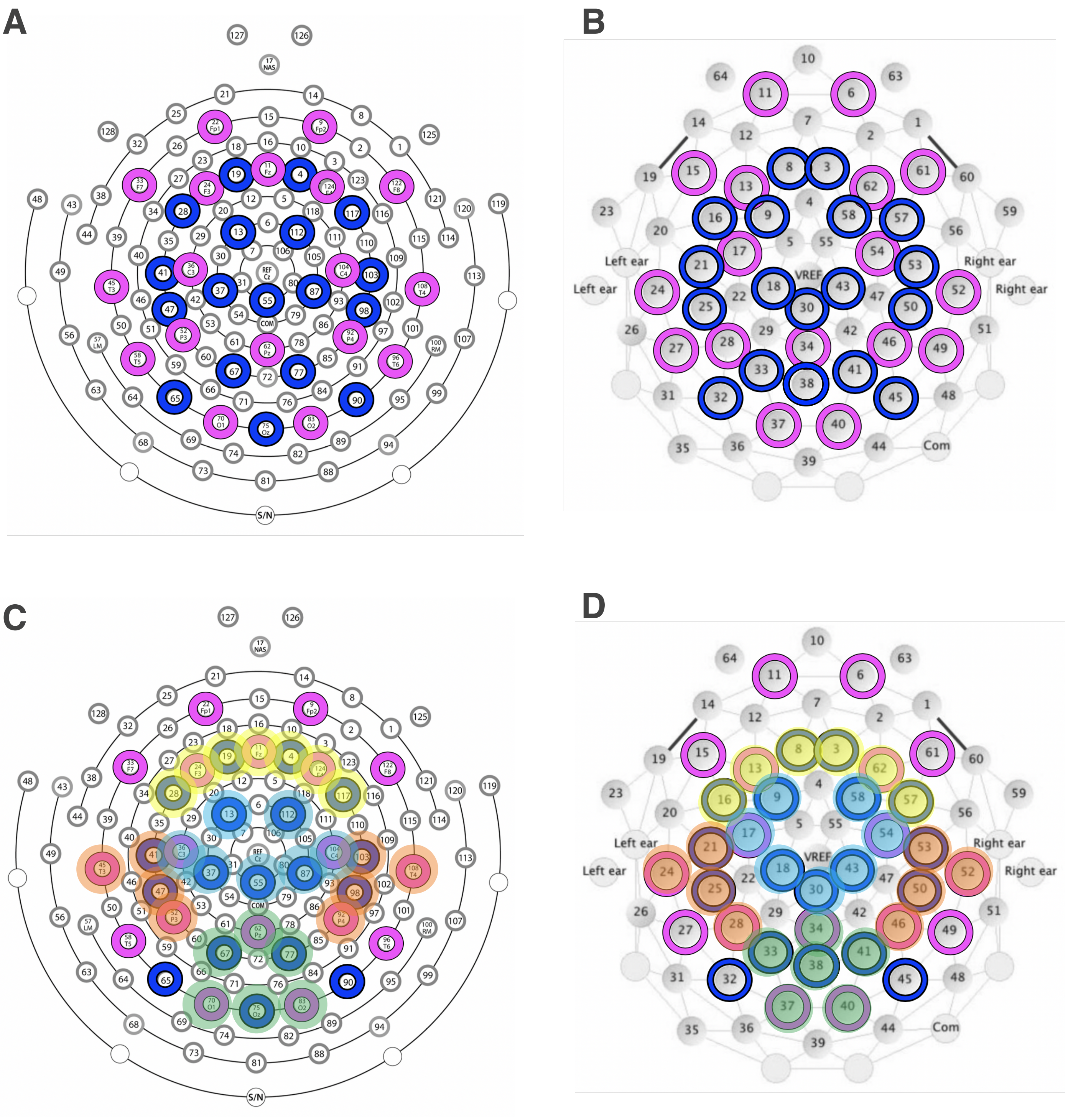

**Supplemental Figure 2:** Individual plots of the periodic spectrum averaged across the whole ROI for each age bin.

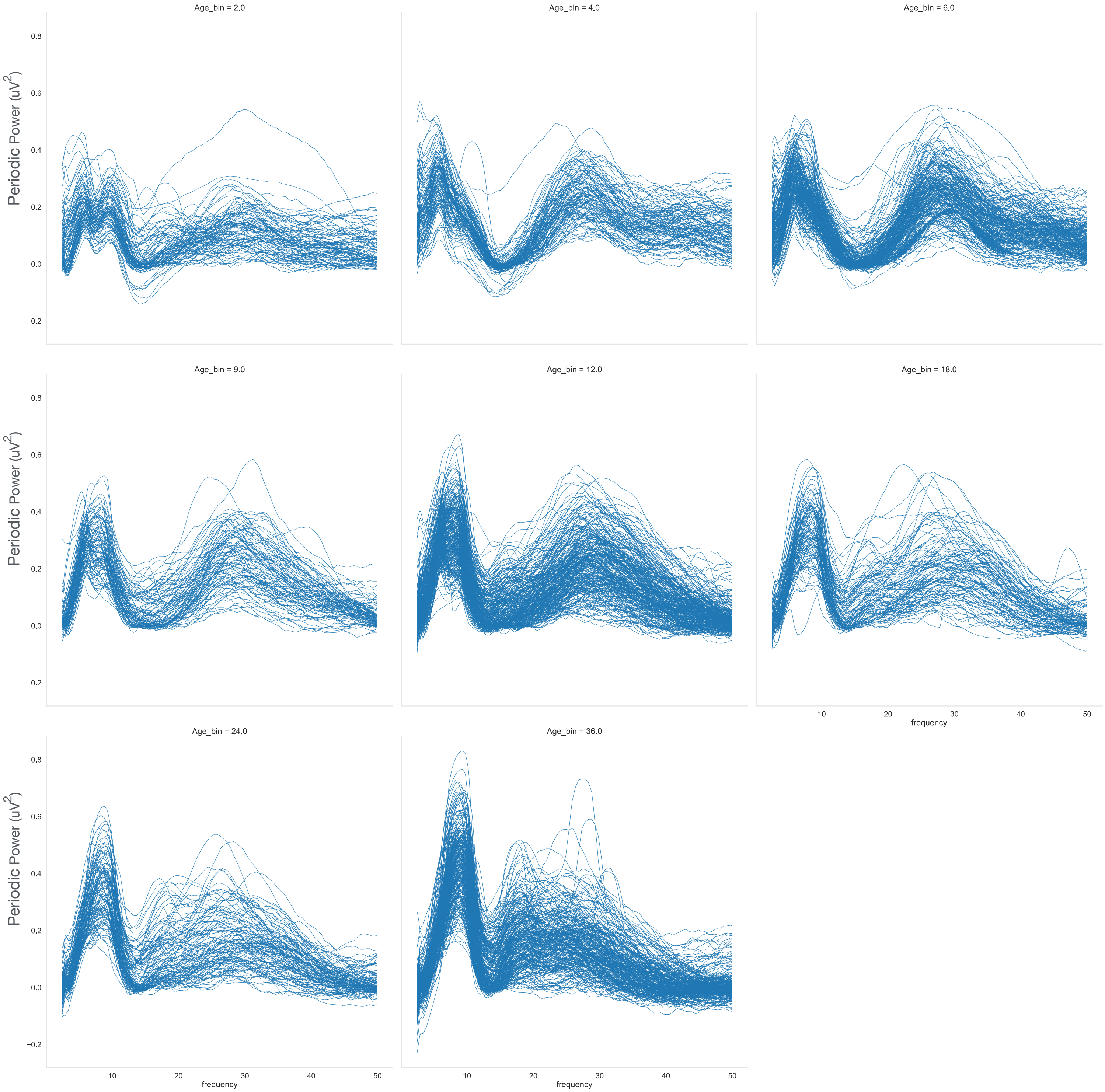

**Supplemental Figure 3:** Spaghetti plots of aperiodic and periodic power measures

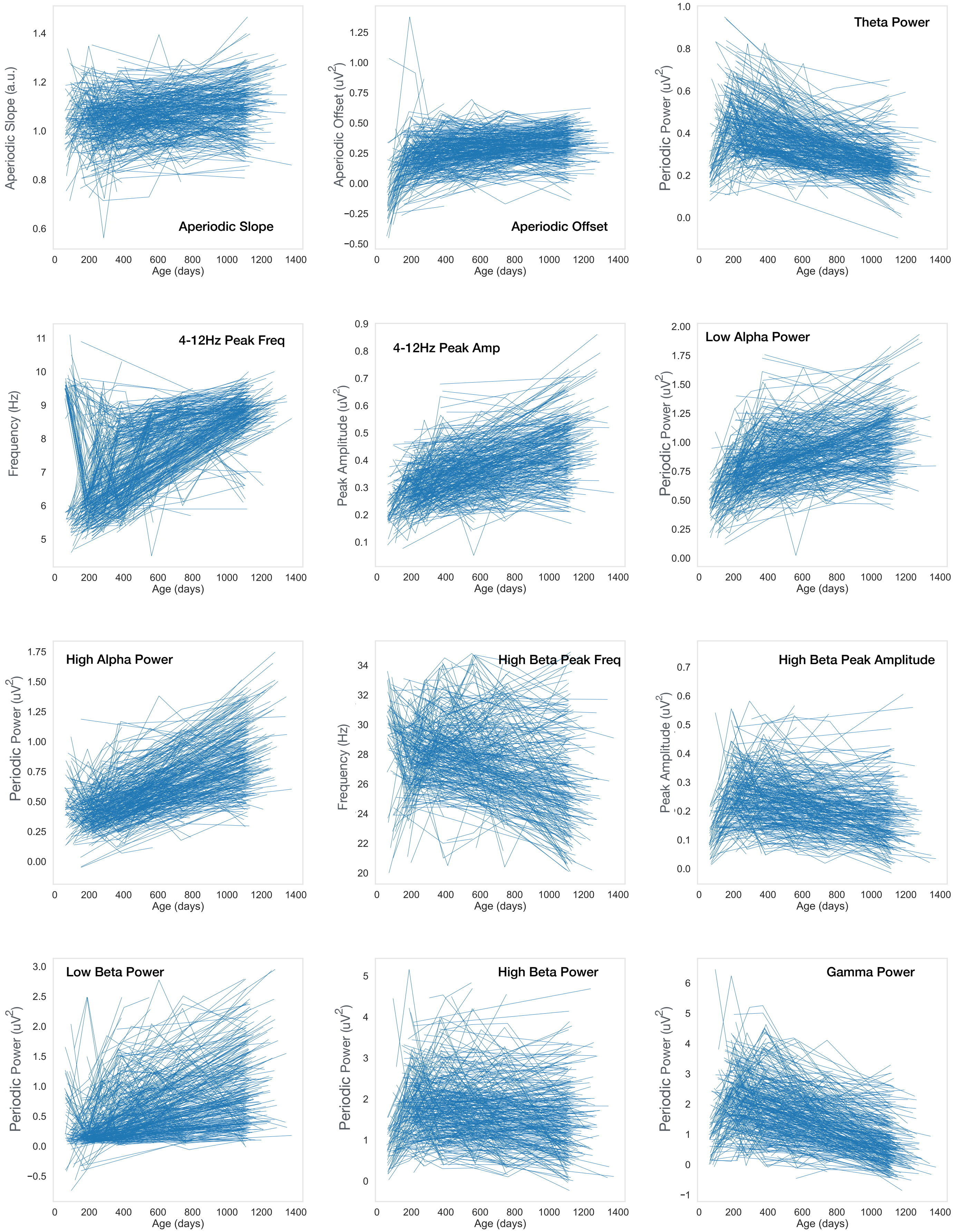

**Supplemental Figure 4:** Spaghetti plots of aperiodic and periodic power measures limited to participants with more than 4 data points and graphed by Age bins.

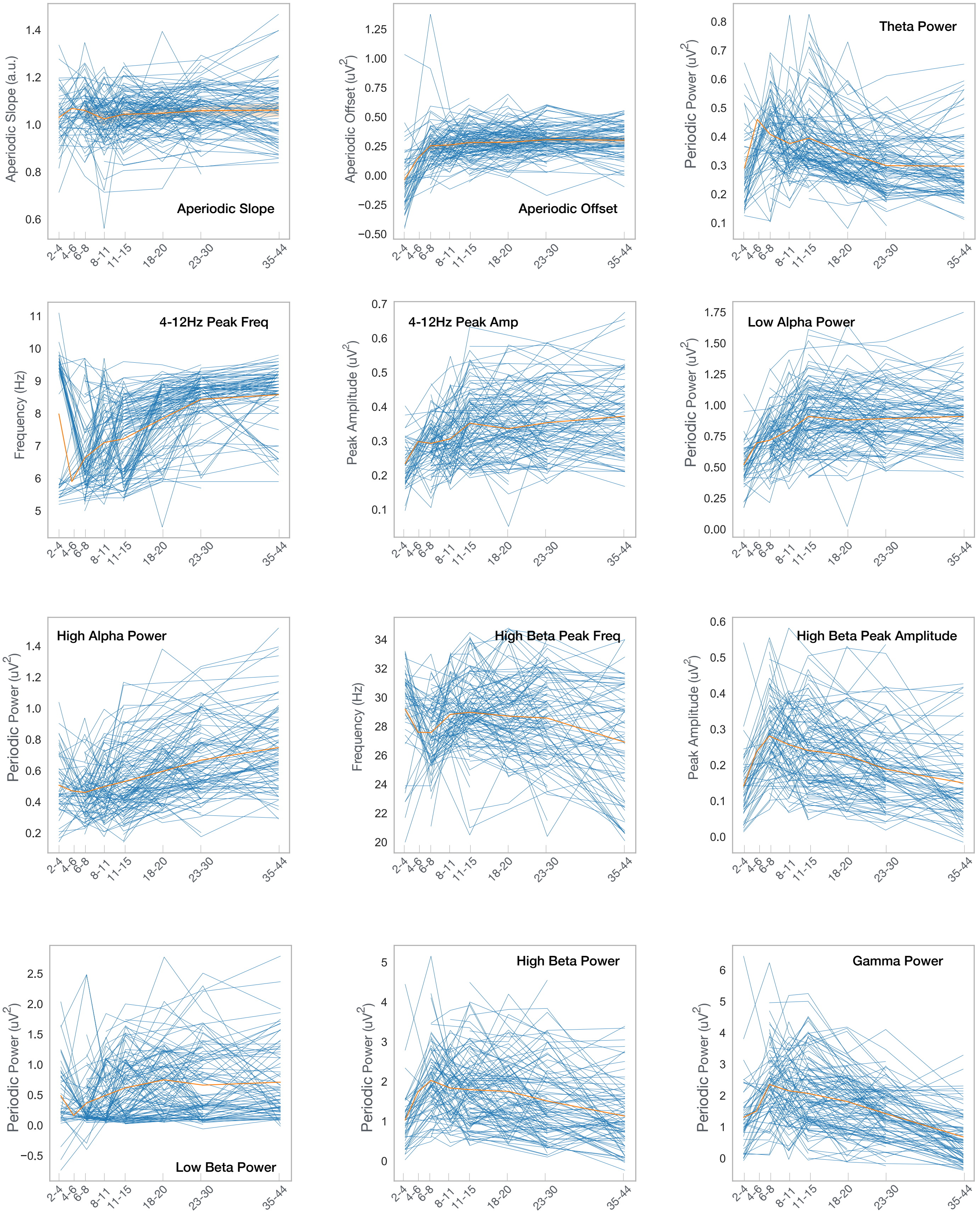

**Supplemental Figure 5:** GAMMs modeled trajectories of 12 aperiodic and periodic power measures, with ROI, study, smoothed age, and sex as predictor terms.

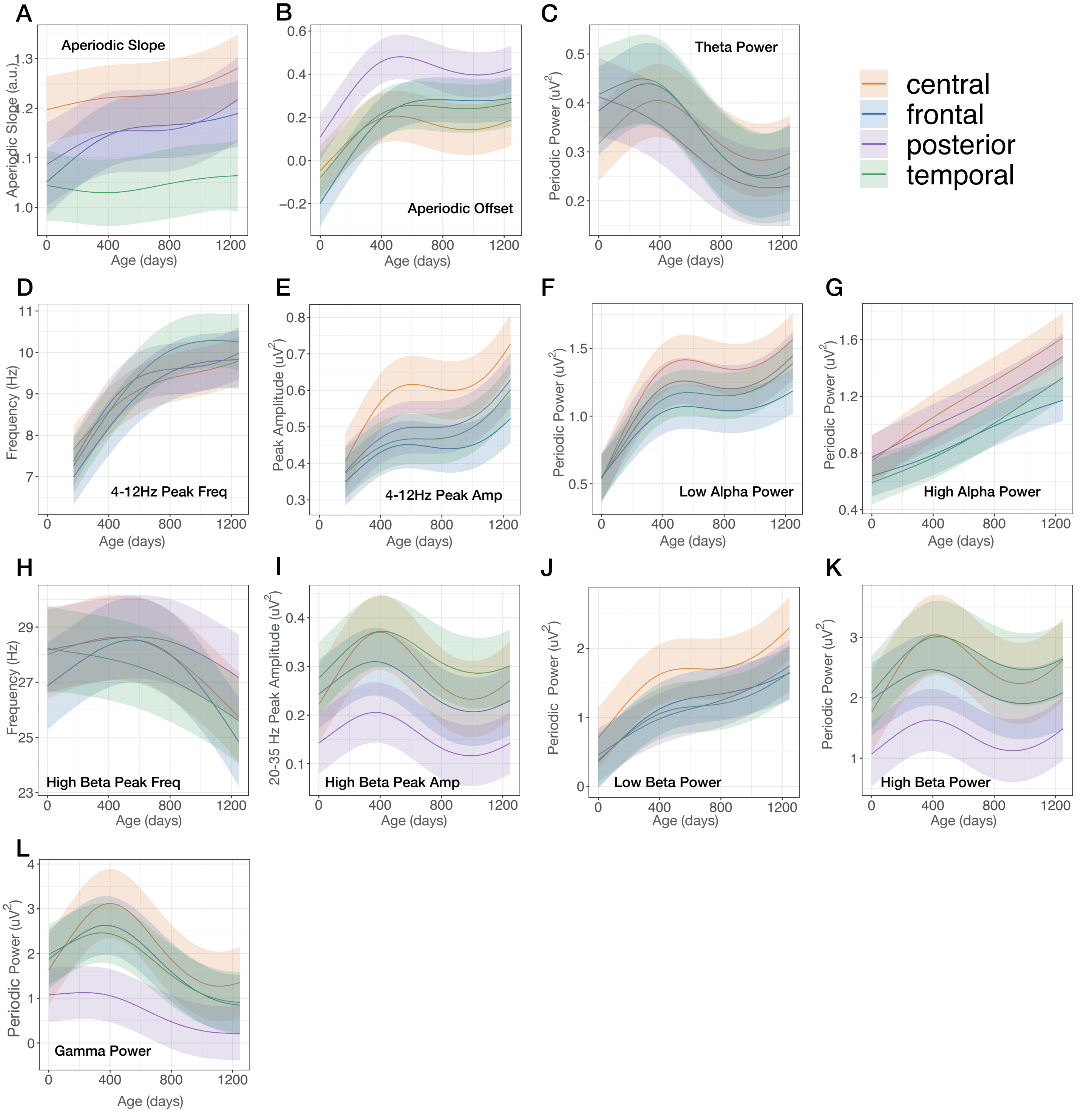

### Supplemental Figure 6

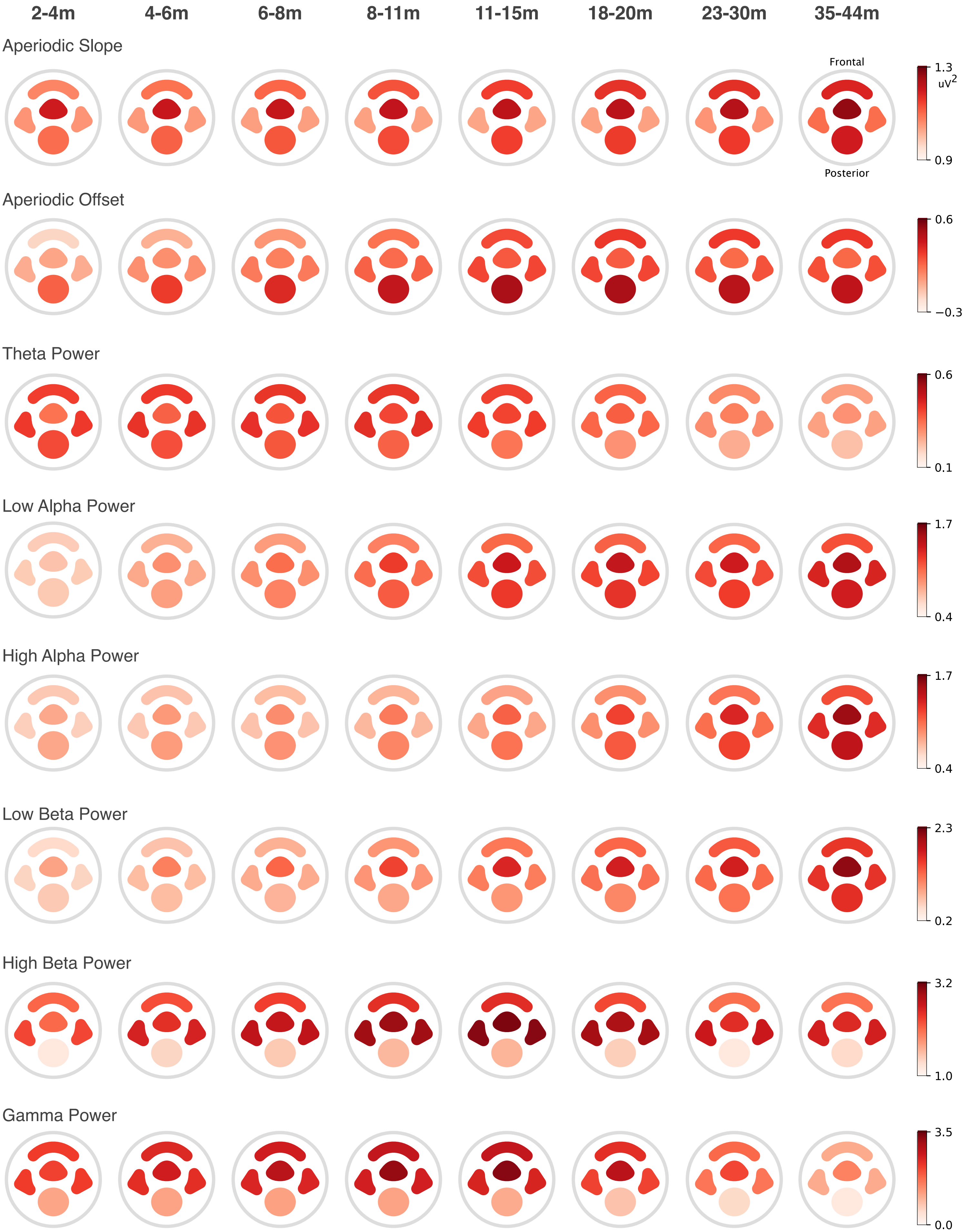

### Supplemental Figure 6 cont.

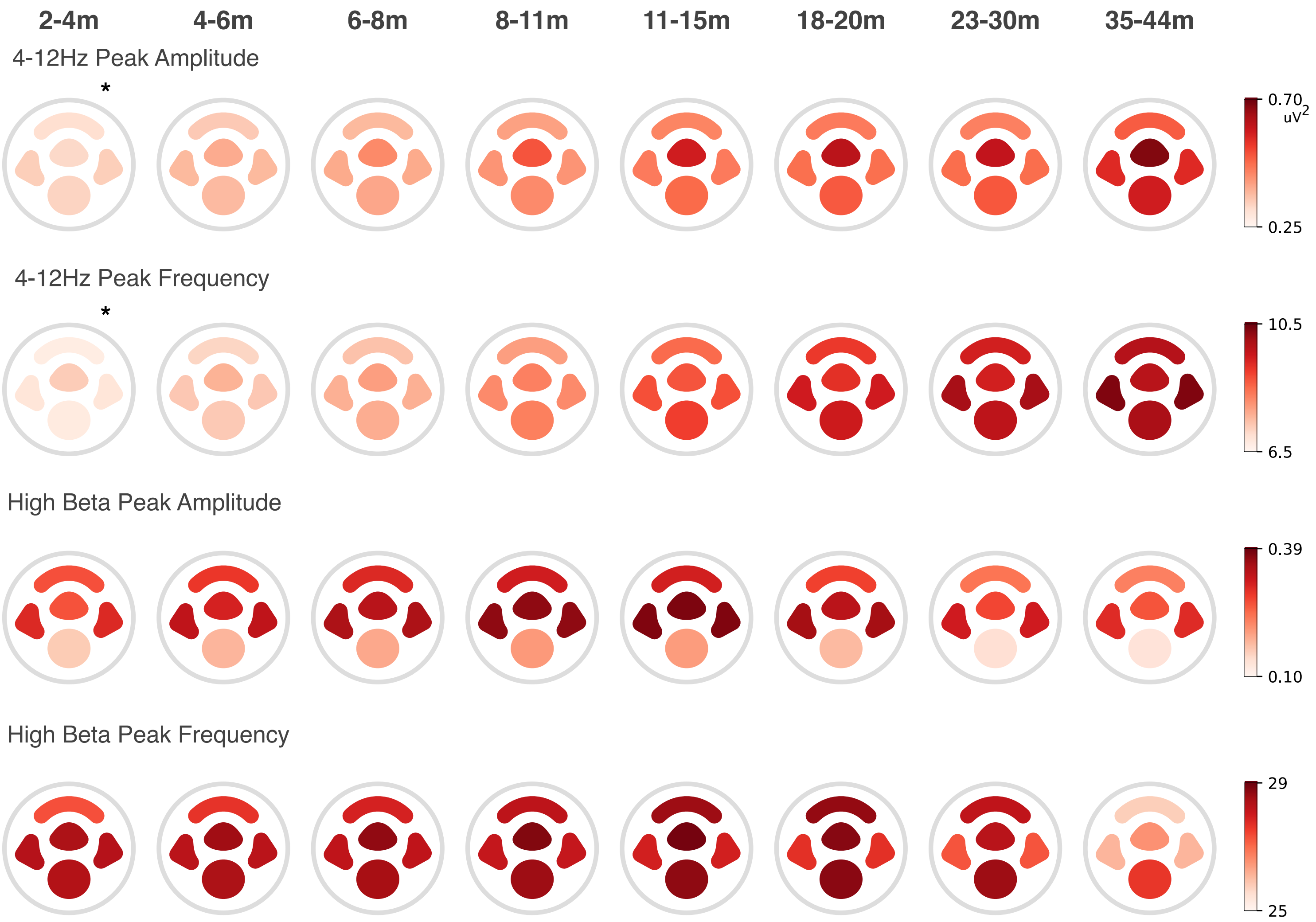


**Supplemental Figure 6:** Modeled estimates of the 12 aperiodic and periodic power measures from the for the four ROIs shown in Supplemental Figure 1 are shown topographically.

*Note, at 2-4 months two peaks are present in the 4-12hz frequency range. Peak amplitude and frequency shown here is based on which ever peak had higher amplitude.

**Supplemental Figure 7:** For each age bin the average SpecParam modeled spectra (green) across participants in each age bin is graphed along with the averaged original power spectrum (red), and averaged SpecParam estimated aperiodic spectrum (blue). (A) Unedited SpecParam estimates. (B) Modified SpecParam estimates. (C) Comparison of squared error across frequencies of unedited (orange) and modified (blue) SpecParam.

**A. Unedited SpecParam model estimates** - Original Spectrum - SpecParam modeled Spectrum - Aperiodic Fit

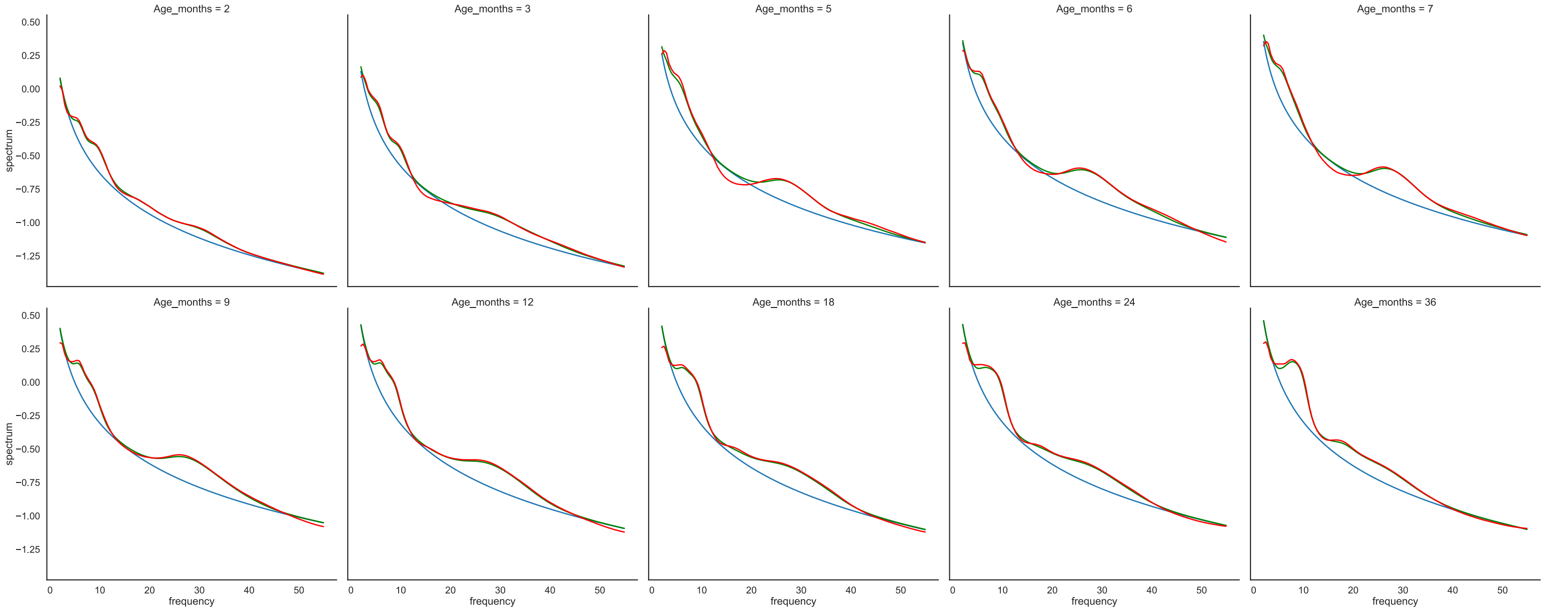


**B. Modified SpecParam model estimates** - Original Spectrum - SpecParam modeled Spectrum - Aperiodic Fit

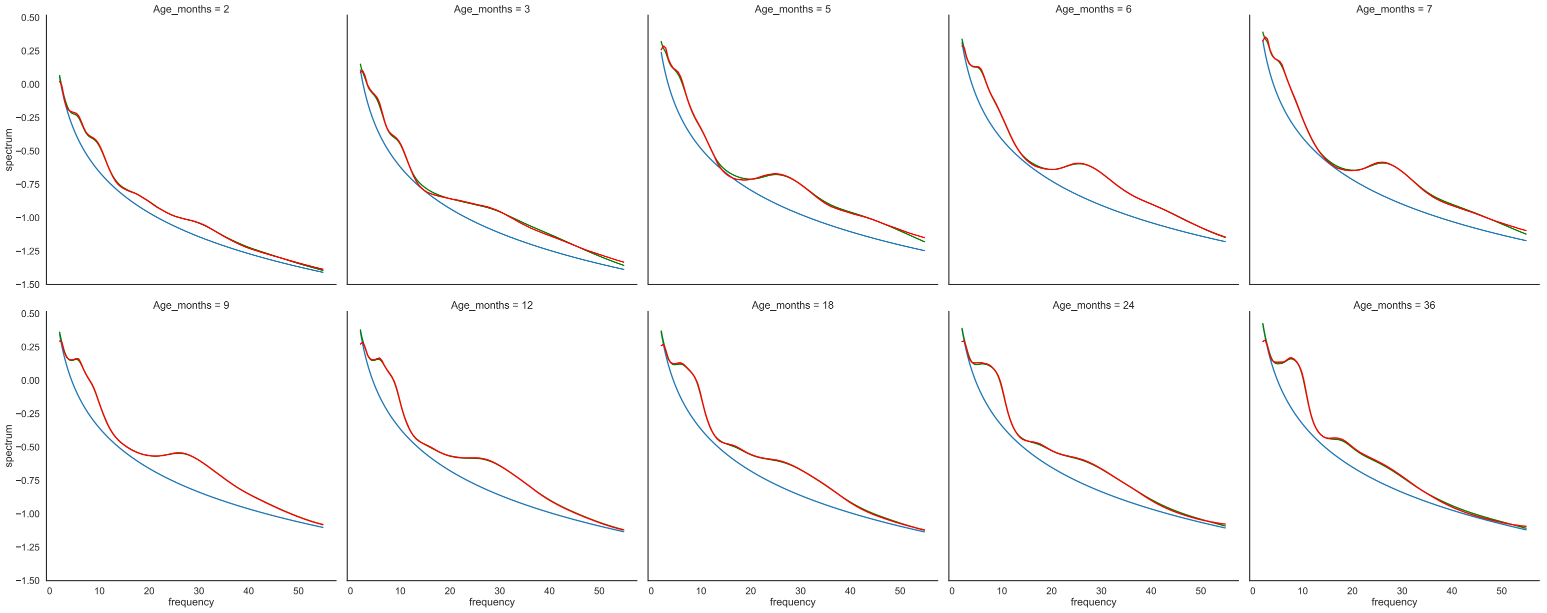

**Supplemental Figure 8:** Comparison of squared error across frequencies based of unedited (orange) and modified (blue) SpecParam estimates of whole brain power.

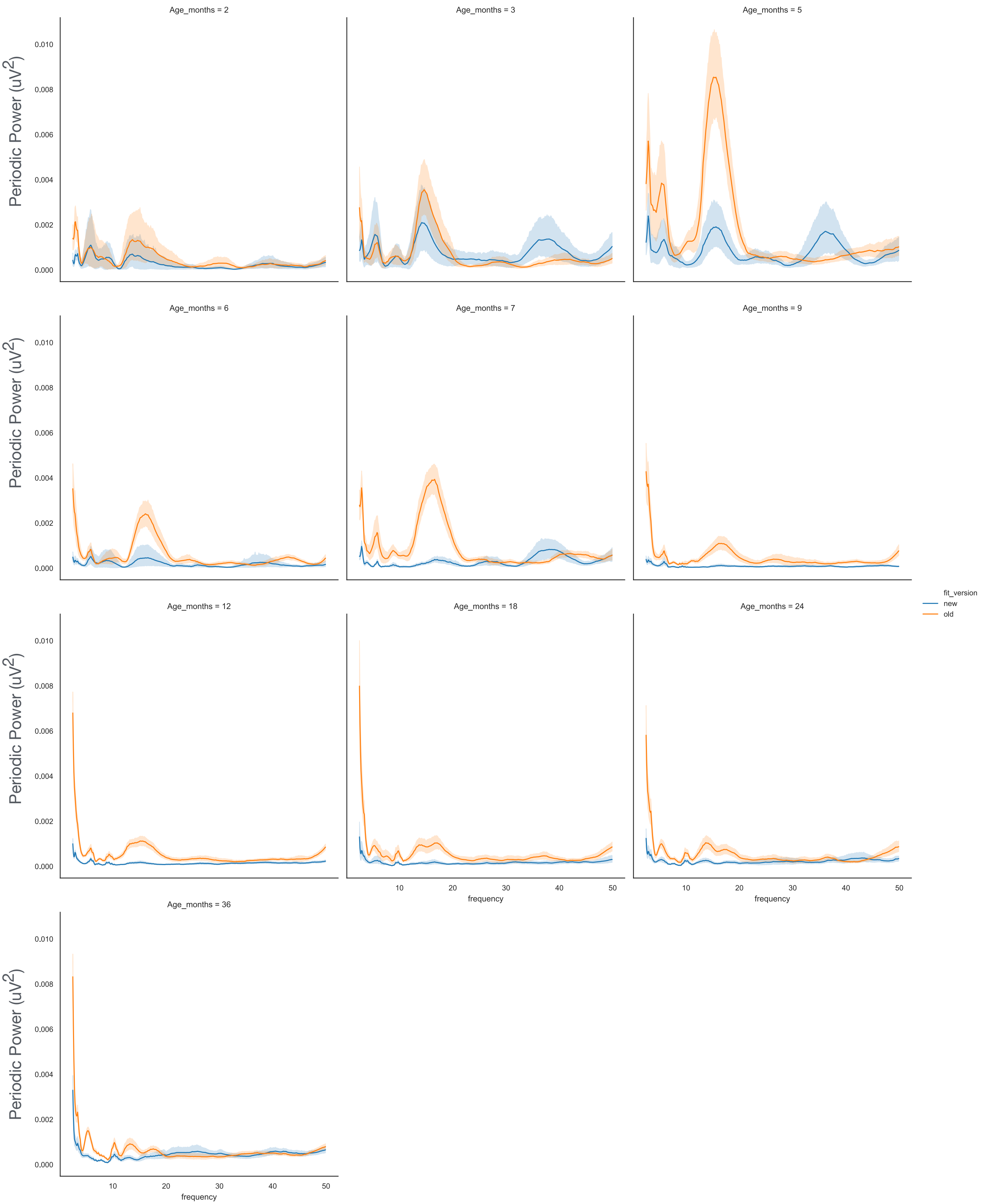

**Supplemental Figure 9:** Periodic power graphed by behavioral scores provided during EEG acquisition of Study 4. No other studies included behavioral ratings. At the time of acquisition a behavioral score 1-5 was provided by the research assistant based on the infant's behavioral regulation during baseline EEG acquisition, with higher scores indicating more behavioral compliance.

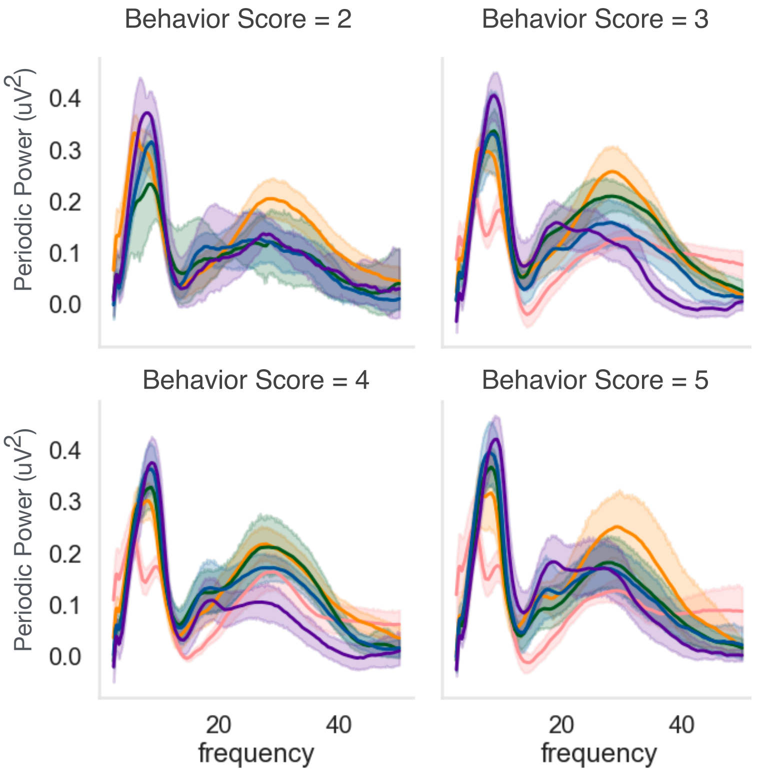


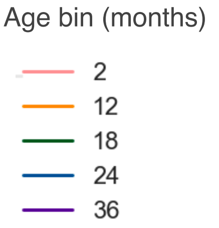


Number of participants

| Behavior Score | | 2m | 12m | 18m | 24m | 36m |
|---|---|---|---|---|---|---|
| 2 | Frequently upset, only short periods of calm between agitation | 0 | 44 | 28 | 22 | 14 |
| 3 | Sometimes upset (30-50% of session) with long periods of calm between protests | 13 | 69 | 42 | 25 | 29 |
| 4 | Rarely upset with agitation/upset 10-30% of session | 17 | 54 | 39 | 33 | 24 |
| 5 | Calm for majority (90%) of session | 10 | 22 | 27 | 18 | 23 |

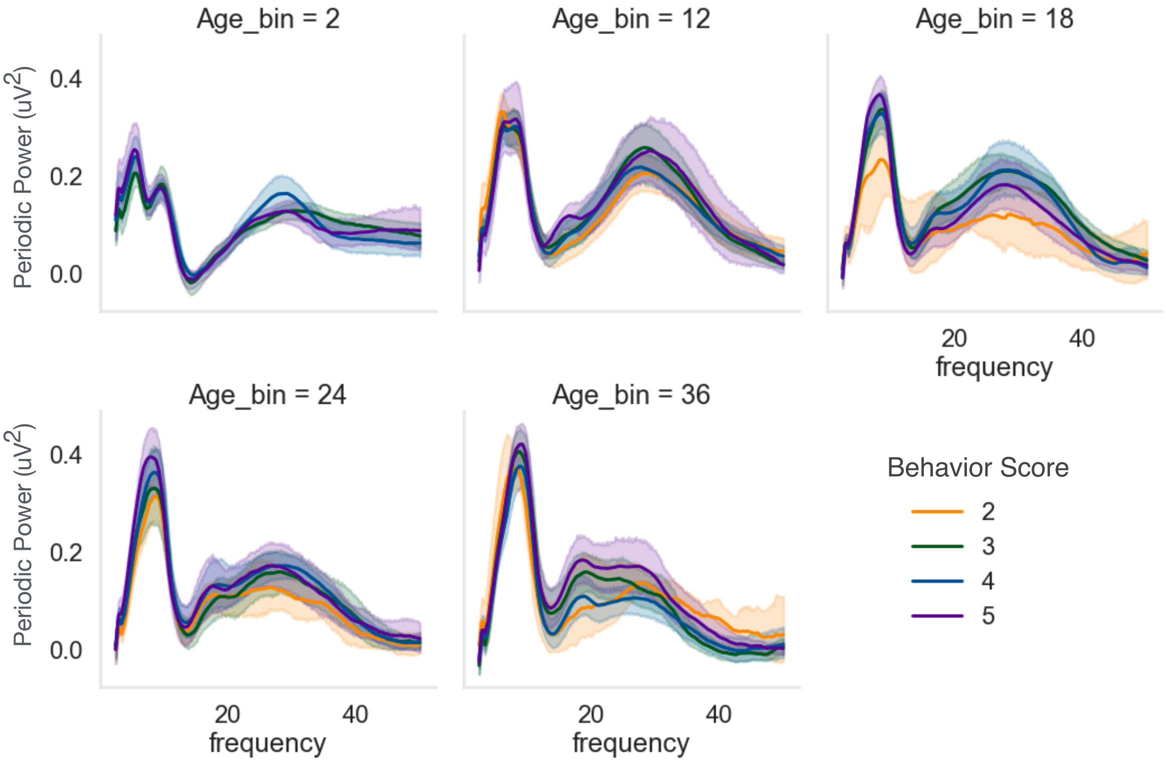

Supplemental Table 1: GAMMs statistics for models of aperiodic offset and slope, and periodic power features

| | Aperiodic Offset | Aperiodic Slope | 4-12Hz Peak Amplitude | 4-12Hz Peak Frequency | High Beta Peak Amplitude | High Beta Peak Frequency |
|---|---|---|---|---|---|---|
| Age F (q value) | 93.42 (0) | 8.43 (0) | 138.3 (0) | 416.62 (0) | 59.31 (0) | 48.59 (0) |
| Sex F (q value) | -0.03 (0.98) | 0.7 (0.60) | -0.54 (0.6708) | 2.23 (0.14) | 1.31 (0.41) | 2.3 (0.14) |
| AgexFemale F (qvalue) | 5.28 (0.0019) | 3.04 (0.0282) | NA (NA) | NA (NA) | NA (NA) | NA (NA) |

Supplemental Table 2: GAMMs statistics for models of periodic power across 6 frequency bands.

| | **Periodic Power** | | | | | |
|---|---|---|---|---|---|---|
| | Theta | Low Alpha | High Alpha | Low Beta | High Beta | Gamma |
| Age F (q value) | 115.12 (0) | 243.83 (0) | 440.63 (0) | 133.04 (0) | 28.19 (0) | 220.69 (0) |
| Sex F (q value) | --1.86 (0.25) | 0.76 (0.60) | 1.27 (0.41) | 0.92 (0.57) | 1.01 (0.56) | 2.9 (0.06) |
| AgexFemale F (q value) | 5.5 (0.0019) | NA (NA) | NA (NA) | NA (NA) | NA (NA) | NA (NA) |

Supplemental Table 3: GAMM model statistics assessing differences between regions of interest.

| | **Periodic Power** | | | | | |
|---|---|---|---|---|---|---|
| | Theta | Low Alpha | High Alpha | Low Beta | High Beta | Gamma |
| Frontal F (q value) | 2.25 (0.025) | -10.61 (0) | -18.45 (0) | 6.88 (0) | 29.76 (0) | 20.1 (0) |
| Central F (q value) | 4.32 (0) | 16.31 (0) | 10.64 (0) | 26.78 (0) | 40.34 (0) | 28.66 (0) |
| Temporal F (q value) | 8.96 (0) | -6.43 (0) | -15.98 (0) | 6.73 (0) | 30.32 (0) | 17.85 (0) |

**Supplemental Methods:**

**Description of code edits to SpecParam. Please see code on Open Science Framework: osf.io/u3gp4**

In the original robust_ap_fit function (https://github.com/fooof-tools/fooof/blob/5e655d73c9d7a47d0411b5177657aaed67c69d6d/specparam/objs/fit.py#L964-L1024), the first estimated flatspec was calculated by subtracting the initial ap fit from the power spectra, AND any value below 0 was converted to 0 (line 993). This leads to the increased error between the original spectrum and the foood estimated spectrum in the 10-20Hz region for 2-7 month olds, since many values fell below 0. Our modified code 'new_robust_ap_fit' elevates the first estimated flat spec such that the lowest point in the flatspec is ≥ 0.

```
    # Flatten power_spectrum based on initial aperiodic fit
    flatspec = power_spectrum - initial_fit

    # OLD: Flatten outliers, defined as any points that drop below 0
    # flatspec[flatspec < 0] = 0  #ORIGINAL
    # NEW: Increase baseline to prevent fitting negative values
    if min(flatspec) < 0:
        flatspec -= min(flatspec)
```

Following this step, a second more robust aperiodic fit is estimated, and then the fit function re-estimates the flattened spectra (spectrum_flat). The original fit function found here: https://github.com/fooof-tools/fooof/blob/5e655d73c9d7a47d0411b5177657aaed67c69d6d/specparam/objs/fit.py#L427-L525. Our modified code then sets any negative data in the flattened spectra equal to 0 (similar to the approach used in the original code during the initial aperiodic fit).

```
    # In rare cases, the model fails to fit, and so uses try / except
    try:

        # Fit the aperiodic component
#         self.aperiodic_params_ = self._robust_ap_fit(self.freqs, self.power_spectrum)  #ORIGINAL
        self.aperiodic_params_ = self._new_robust_ap_fit(self.freqs, self.power_spectrum)  #NEW
        self._ap_fit = fooof.sim.gen.gen_aperiodic(self.freqs, self.aperiodic_params_)  #edited to make standalo

        # Flatten the power spectrum using fit aperiodic fit
        self._spectrum_flat = self.power_spectrum - self._ap_fit

        self._spectrum_flat[self._spectrum_flat < 0] = 0  #NEW
```
